# Human EEG Decoding Reveals Lapse-Prone States Beyond Selective Attention and Control Failures

**DOI:** 10.1101/2025.09.24.678372

**Authors:** Matthieu Chidharom, Henry M. Jones, Monica D. Rosenberg, Edward K. Vogel

## Abstract

Attentional lapses are a ubiquitous feature of cognition, yet their underlying causes remain poorly understood. Theories of sustained attention often point to failures of cognitive control in maintaining the task-set, while data-driven approaches suggest that lapses may instead reflect a breakdown in the selection of task-relevant information. This study aimed to characterize the neural mechanisms of sustained attention lapses and to test whether EEG-based signatures of lapse-prone states are distinct from signatures of failures of selective attention and task-set maintenance. Twenty adults completed a sustained attention go/no-go task while focusing on either numbers or letters, with EEG recorded simultaneously. Poor sustained attention was examined at two complementary levels: trial-level lapses, defined as no-go errors, and attentional states, derived from reaction-time variability and categorized as “in-the-zone” versus “out-of-the-zone”. Across both levels, suboptimal sustained attention was associated with attenuated event-related potentials, most notably a reduced parietal P3 amplitude and weaker whole-scalp inter-electrode correlation. To isolate a unique EEG marker of lapse-prone state, a machine-learning classifier decoded attentional state from EEG activity. Cross-validated accuracy reached ∼80% and remained robust after controlling for reaction time. Finally, representational similarity analysis confirmed that this neural signature was dissociable from stimulus-side selection and task-set maintenance.

## Introduction

Lapses of sustained attention occur when attention drifts away from the task at hand ^1^. In today’s hyperconnected world, these lapses feel increasingly common, with serious consequences for productivity, learning, and safety. But why do lapses occur? Is it because we fail to select what matters—like checking our phone instead of reading? Is it because we lose track of our goals and forget what we’re doing? Or are lapse-prone states driven by something else entirely? This study aims to better characterize the neural basis of sustained attentional lapses and to predict their occurrence using machine learning approaches.

In cognitive psychology, the capacity to resist distractions and to maintain focus over time is known as sustained attention. When this ability fails, we experience attentional lapses—brief moments of attentional disengagement. These lapses have become central to recent research on concentration. To investigate them, researchers have long relied on go/no-go or continuous performance tasks (CPTs). In these paradigms, participants are instructed to respond to frequent “go” stimuli (e.g., digit numbers) and to withhold responses to rare “no-go” stimuli (e.g., the digit number 3), which typically occur on 10% of trials ^2^. Failures to withhold a response on no-go trials are taken as markers of attentional lapses, and have been associated with everyday distractions in work and life ^2–5^.

To predict when lapses occur, researchers have adopted two main approaches: theory-driven and data-driven. Within the theory-driven perspective, there is broad agreement that cognitive control plays a central role in maintaining task goals and resisting interference. Several influential frameworks, including the control failure hypothesis ^6^, the oscillatory model ^7^, and the resource-control account ^8^, converge on the idea that lapses arise when cognitive control fails to sustain the task-set over time. Supporting evidence comes from studies on inter-individual differences, showing that individuals with weaker cognitive control are more susceptible to lapsing ^9,10^, as well as from neuroscience findings linking lapses to decreased neural markers of control ^7,11–15^. However, these studies mostly use sustained attention tasks in which only one stimulus is presented at the center of the screen. Consequently, this feature makes it difficult to determine whether a failure of cognitive control is the only cause of attention lapses, or if impaired selective attention could also explain lapses, as suggested by Weissman and colleagues ^16^ and more recent data-driven studies.

The data-driven approach aims to predict lapse likelihood based on behavioral patterns or neural activity. On the behavioral side, major advances have been made. Esterman and colleagues ^17^, using the Variance Time Course (VTC) method, identified periods of high response variability—termed “out-of-the-zone”—and periods of low variability—“in-the-zone.” They showed that out-of-the-zone periods predicted upcoming no-go errors, suggesting that increased behavioral variability marks moments of lapse-prone attentional states. Another behavioral method proposed by deBettencourt and colleagues ^18^ focused on reaction times and showed that short RTs uniquely predict upcoming lapses in CPTs.

On the neural side, progress in predicting lapse from fMRI data has been consistent, whereas findings based on EEG remain more heterogeneous and modest. For instance, Kaushik and colleagues ^19^ applied machine learning techniques to classify lapse-prone states using EEG recordings in a real-world setting. However, these classifications relied on subjective reports, as the ecological context made it difficult to isolate attentional fluctuations objectively. Similarly, a recent study attempted to decode rare omission errors using EEG in a controlled task, but predictive accuracy remained limited ^20^. Complementary advances have been made using fMRI. Rosenberg and colleagues used functional connectivity analyses to identify high- and low-attention networks that predict lapse-proneness at the individual ^21^ and state level ^22^. At the intra-individual level, work has also identified distinct brain states associated with lapse frequency during CPTs: one dominated by activity in the default mode network was linked to better concentration, whereas another characterized by greater engagement of the dorsal attention network (DAN) was associated with more frequent lapses ^23,24^.

These data-driven results provide new insights that refine our understanding of how lapses occur. Among these findings, the increase in DAN activity during out-of-the-zone states has been replicated several times, and DAN disengagement has been observed just before the occurrence of no-go errors (or lapses) ^17,25,26^. These results suggest that during lapse-prone states, individuals allocate more effort in processing task-relevant stimuli and that lapses arise when attention fails to select the relevant information.

Taken together, theoretical accounts suggest that sustained attention lapses are predicted by a failure of cognitive control to maintain the task-set in mind, while recent data-driven results argue in favor of an impaired selective attention to relevant information. A question therefore arises: is predicting a lapse simply decoding the efficiency of cognitive control and selective attention, or are lapses associated with distinct and unique neural signals, independent of other forms of attention?

To answer this question, we engaged participants in a bilateral CPT. Participants were instructed to perform a go/no-go task on either the letters or the numbers, which were presented simultaneously on the left or right side of a fixation point. This task design allowed us to investigate how cognitive control maintains the task-set in mind (numbers vs. letters) and how selective attention allocates resources to the task-relevant side (left vs. right).

As a first step, we aimed to better characterize the EEG components altered during poor sustained attention. We identified a lack of concentration at the trial level by focusing on no-go errors in the CPT and at the state level by isolating in-the-zone and out-of-the-zone periods ^17^. Second, we sought to test whether lapse-prone states could be predicted and isolated from EEG signals beyond other attentional functions or behavioral measures. To this end, we employed machine learning approaches using multivariate classification and representational similarity analysis (RSA), which allowed us to isolate a unique EEG signature of sustained attention states with an accuracy of 80%, that is independent of other forms of cognitive control and selective attention.

## Results

To study poor sustained attention, we used a bilateral-CPT, in which 20 participants viewed a number and a letter presented either to the left or right of a central fixation point **(Figure 1A)**. Every 20 trials, a cue appeared on the screen, randomly indicating which task the participant should perform. In the number task, participants were instructed to press a response button as quickly as possible whenever a digit appeared (go trial), except when the digit was 3 (no-go trial). In the letter task, participants had to press the button in response to any letter except the letter K. The task consisted of 3,200 trials and lasted approximately two hours. On average, 24% of trials (SD = 15.1) were rejected for artefacts, leaving us with a reliable dataset of approximately 48,500 trials. Using this task, our first goal was to examine how poor sustained attention—at the trial- and state-level—modulates neurophysiological markers of attention. Second, we tested whether lapse-prone states could be predicted directly from EEG signals using a machine learning approach, and whether this neural signature is unique—or merely reflects other known forms of attention, such as selective attention or cognitive control.

**Figure 1.**
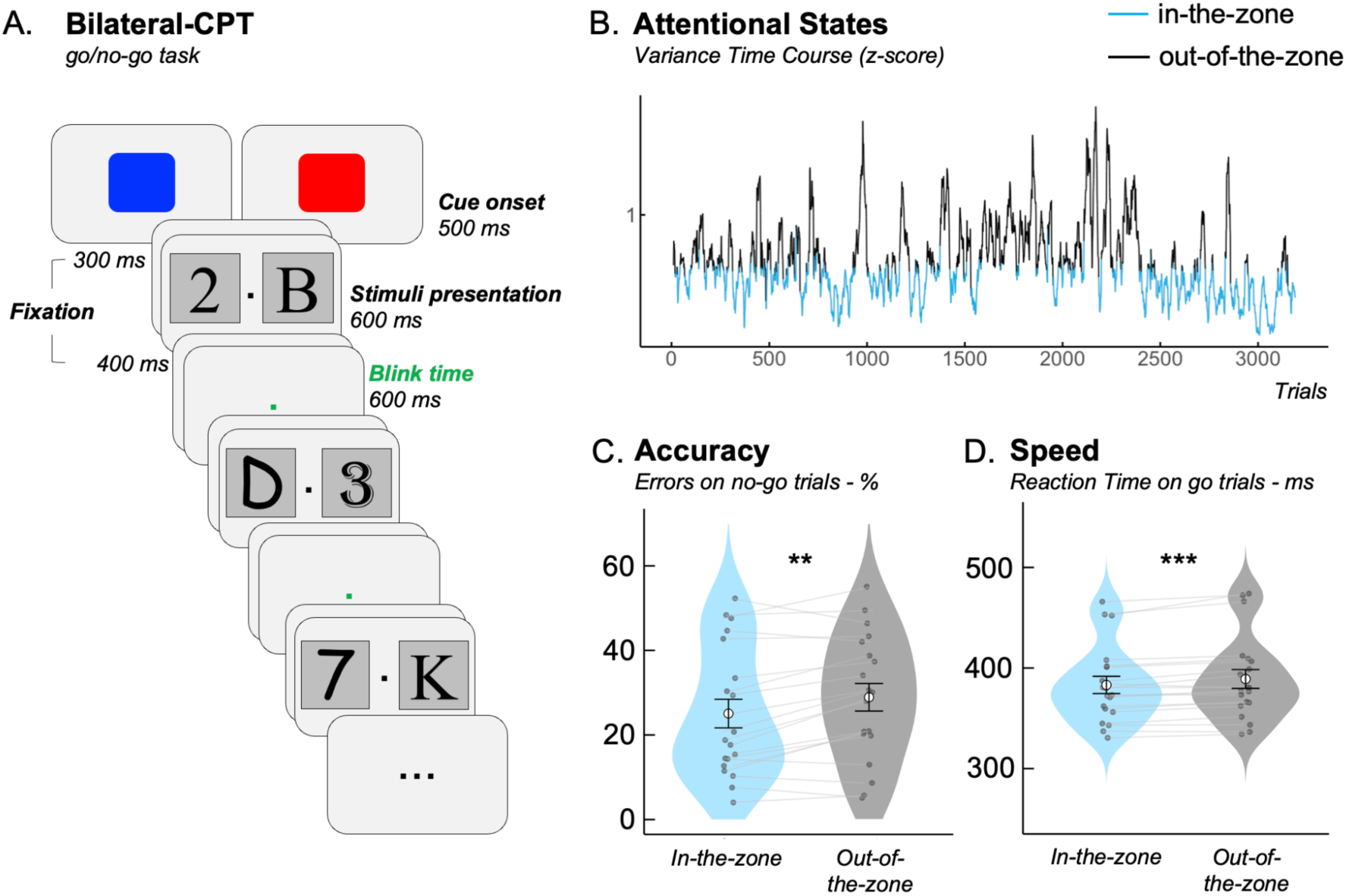
Task and behavioral performance. **A.** Bilateral CPT. Participants were instructed to focus on either letters or numbers depending on a cue presented every 20 trials. For example, when the cue was blue, participants responded to all numbers except the digit 3 (no-go). When the cue was red, they responded to all letters except the letter K. **B.** VTC calculation. Example for one participant. The Variance Time Course (VTC) is shown across 3,200 trials, highlighting the identification of in-the-zone and out-of-the-zone periods. The VTC has been smoothed for illustrative purposes. **C.** Accuracy. The VTC method effectively isolated periods of poor sustained attention: participants made significantly more lapses (i.e., no-go errors) during out-of-the-zone than in-the-zone periods. **D.** Reaction time. Reaction times on correct go trials were significantly slower during out-of-the-zone compared to in-the-zone periods. \*\**p* < .01; \*\*\**p* < .001.

### Performance

Reaction times (RTs) and error rates on no-go trials closely mirrored those reported in prior research, with participants responding rapidly (M = 386 ms, SE = 8.95) and making errors on 26.8% of no-go trials (SE = 3.29). Omission rates on go trials remained minimal (M = 1.13%, SE = 0.216). Critically, we successfully replicated a hallmark pattern observed in unilateral CPTs (deBettencourt et al., 2019), whereby reaction times preceding correct no-go were significantly longer (M = 386 ms, SE = 8.32) than those preceding no-go errors (M = 355 ms, SE = 8.52), *t*(19) = 8.50, *p* < .001, *d* = 1.90. This finding suggests that the bilateral CPT produces behavioral effects similar to those observed in the unilateral version. Performance in the letter and number tasks did not differ, either on RT or on no-go errors (*see SI Results*).

### Effect Of Lapses of Sustained Attention on the EEG Signal

Having replicated classic behavioral effects, our first goal was to examine how lapses (i.e., no-go errors) impact neurophysiological markers. To do so, we segmented the EEG signal time-locked to stimulus onset and first focused on the effects of lapses on ERPs. A key component of interest was the parietal P300—a well-established marker of attentional allocation. If attention is indeed withdrawn from the ongoing task during no-go errors, then the P300 should be reduced during these errors, as previously demonstrated in studies on attentional lapses ^14,27^. We replicated this effect in our task, as confirmed by an ANOVA on P300 amplitude that revealed a significant main effect of trial type, F(2,38) = 36.0, p < .001, η²p = .655. Post-hoc t-tests further revealed that P300 amplitude was significantly lower for no-go errors compared to no-go correct trials, t(19) = 3.20, p = .005, p_tukey_ = .012, consistent with a failure to allocate sufficient attentional resources during lapses **(Figure 2A)**. Interestingly, both no-go correct and no-go error trials elicited higher P300 amplitudes than go trials (no-go correct vs. go: t(19) = –8.23, p < .001, p_tukey_ < .001; no-go error vs. go: t(19) = –5.13, p < .001, p_tukey_ < .001), highlighting the inherently greater attentional demand on no-go trials.

**Figure 2.**
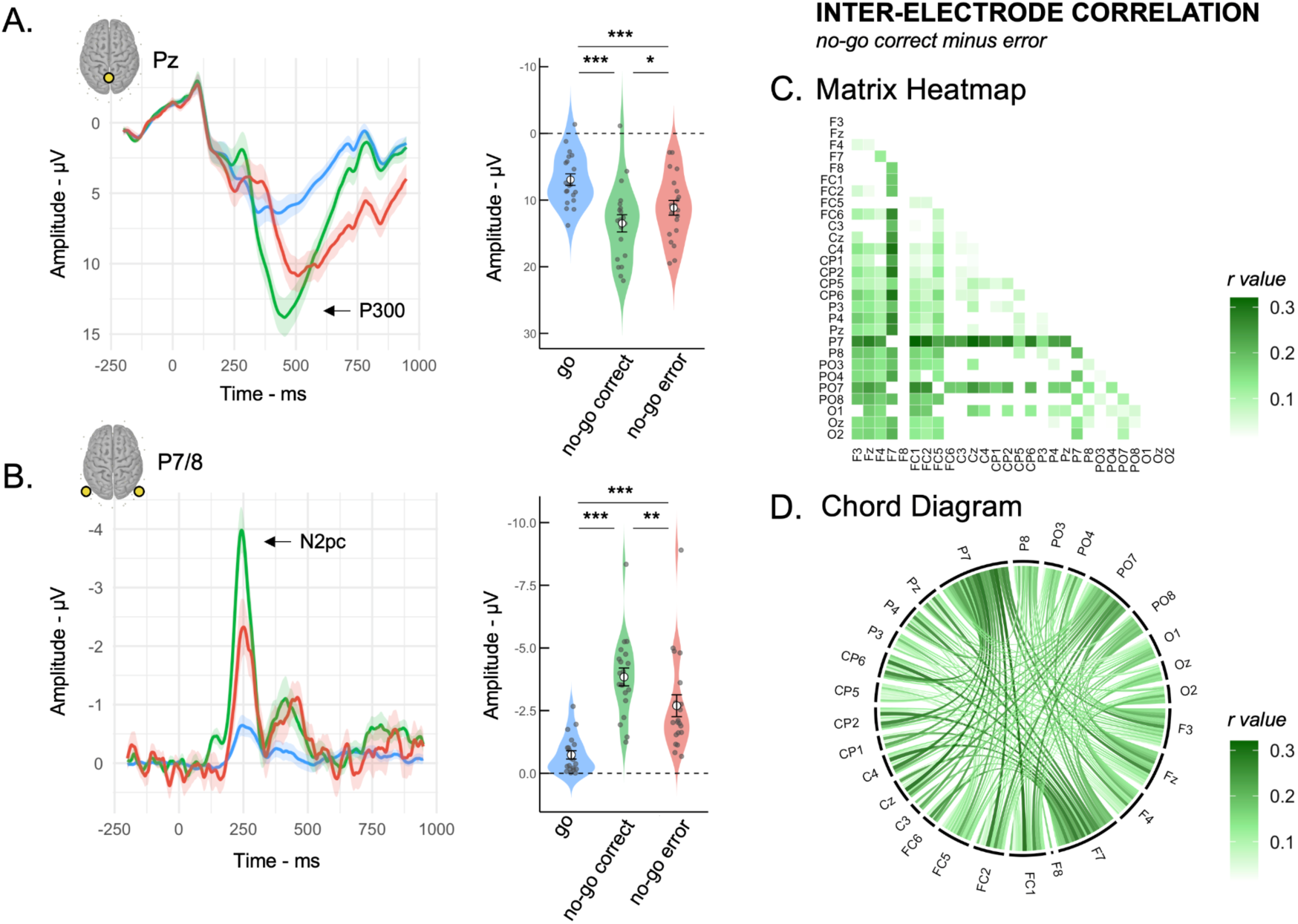
Neural mechanisms of sustained attention lapse. Lapsing was assessed through no-go errors. The amplitudes of the P300 **(A)** and the N2pc **(B)** were reduced during no-go errors compared to correct no-go. Shaded area represents the standard error of the group mean. The difference in inter-electrode correlation scores between correct and error no-go trials revealed increased correlations during periods of high attention, as shown in the FDR-corrected correlation matrix **(C)** and the circular correlation diagram **(D)**. \**p* < .05; \*\**p* < .01; \*\*\**p* < .001.

Beyond the P300, the bilateral CPT offers a unique opportunity to investigate how lapses modulate the spatial allocation of attention toward goal-relevant stimuli—a dimension often overlooked in studies of sustained attention. A key lateralized neural marker is the N2pc, indicating how attention is allocated toward spatially relevant stimuli. Yet, to our knowledge, no prior study has examined how this component behaves during attentional lapses. Our analyses revealed a clear modulation of the N2pc component as a function of trial type, F(2,38) = 48.5, p < .001, η²p = .718 **(Figure 2B)**. Post-hoc comparisons showed that N2pc amplitude was significantly reduced for no-go error trials compared to no-go correct trials, t(19) = –3.94, p < .001, p_tukey_ = .002, suggesting a failure to allocate spatial attention toward the goal-relevant stimuli during lapses. Importantly, N2pc amplitude was higher for both no-go correct, t(19) = 10.26, p < .001, p_tukey_ < .001 and no-go error trials, t(19) = 5.46, p < .001, p_tukey_ < .001, relative to go trials, indicating that no-go stimuli are more effective at capturing spatial attention, irrespective of response accuracy **(Figure 2B)**. To determine whether the reduced N2pc amplitude in error trials was driven by diminished processing of the target or increased distraction by irrelevant stimuli, we conducted a follow-up ANOVA on contralateral and ipsilateral waveforms. The results revealed a significant reduction in contralateral—but not ipsilateral—activity during no-go errors (*see SI Results,* **Figure S1**), suggesting that attentional lapses primarily reflect a failure to select the relevant target, rather than increased attentional capture by irrelevant stimuli.

In addition to ERP analyses, we examined how lapses impact the synchrony of ERP waveforms by analyzing inter-electrode correlations.^7,11,24^. Inspired by methods developed by Hakim and colleagues who found that patterns of inter-electrode correlations better predict individual differences in behavior than individual ERP amplitudes alone, we computed trial-averaged EEG signals for each electrode and calculated the Pearson correlation between every pair of electrodes, separately for correct and error no-go trials ^28^. This yielded whole-scalp inter-electrode correlation matrices for each condition (*see SI Methods*). As shown in **Figures 2C and 2D**, correct no-go trials were associated with stronger inter-electrode correlations compared to no-go errors. Specifically, we observed increased correlations between ERPs during successful inhibition trials. To statistically assess these differences, we computed a difference matrix (no-go correct – no-go error) and applied a false discovery rate (FDR) correction to identify significant correlations (*p* < 0.05). This analysis confirmed a significant enhancement in inter-electrode coupling during correct trials, suggesting that periods of concentration are supported by more globally synchronous trial-evoked neural activity. This result aligns with previous findings that periods of lapse were associated with a reduction in phase oscillatory synchronization compared to periods of stronger concentration ^13,15^.

### Effect Of Lapse-Prone States on the EEG Signal

Having characterized how lapses affect EEG signals at the trial-level, our next goal was to explore how lapse-prone states affect EEG. To do so, we first aimed to identify periods of heightened lapsing based on participants’ behavior and to examine how these periods influenced neural activity. Specifically, we sought to replicate the Variance Time Course (VTC) method introduced by Esterman et al. (2013), which classifies time windows as “in-the-zone” (low RT variability) or “out-of-the-zone” (high RT variability) **(Figure 1B)** (*see SI Methods*). This approach successfully isolated periods of increased lapsing in our task: participants made significantly more no-go errors during out-of-the-zone periods (M = 28.93%, SE = 3.27) compared to in-the-zone periods (M = 25.07%, SE = 3.38), *t*(19) = –3.50, *p* = .002, *d* = –0.78 **(Figure 1C)**. Reaction times on correct go trials were also slower out-of-the-zone (M = 389, SE = 9.34) compared to in-the-zone periods (M = 383, SE = 8.57), t(19) = –4.53, *p* < .001, *d* = –1.01 **(Figure 1D)**.

We next examined how behavioral states (in-the-zone vs. out-of-the-zone) influenced both ERP markers and inter-electrode correlations. For the P300 component, a repeated-measures ANOVA revealed a significant effect of state, with reduced amplitude during out-of-the-zone periods (M = 10.4 µV, SE = 1.02) compared to in-the-zone periods (M = 10.9 µV, SE = 0.97), *F*(1, 19) = 6.23, *p* = .022, η²p = .247 **(Figure S2A)**. This finding confirms that out-of-the-zone periods are associated with reduced attentional engagement. The ANOVA also revealed a significant interaction between trial type and attentional state on P300 amplitude, *F*(2,38) = 3.28, *p* = .049, η²p = .147; where the P300 amplitude was only modulated by states on go trials. Indeed, post-hoc comparisons revealed that the P300 amplitude on go correct trials was significantly lower in out-of-the-zone periods (M = 6.40, SE = 0.86) than in-the-zone periods (M = 7.68, SE = 0.88), *t*(19) = 5.66, *p* < .001, p_tukey_ < .001. For no-go correct trials, there was no significant difference between states (M_in = 13.41, M_out = 13.52), *t*(19) = –0.41, *p* = .690, p_tukey_ = .998. For no-go error trials, P300 amplitude also did not differ between states (M_in = 11.54, M_out = 11.19), *t*(19) = 0.63, *p* = .535, p_tukey_ = .987). However, we observed that P300 amplitude was significantly reduced during no-go error trials compared to no-go correct trials—but only in the out-of-the-zone condition, *t*(19) = 3.36, *p* = .003, p_tukey_ = .033; and not in-the-zone: *t*(19) = –2.12, *p* = .047, p_tukey_ = .317. These findings suggest that lapse-prone states are associated with a marked decrease in attentional engagement during critical moments of the task.

Turning to the N2pc component, the ANOVA revealed no significant main effect of attentional state, *F*(1,19) = 0.18, *p* = .680, η²p = .009. N2pc amplitude was comparable between in-the-zone periods (*M* = –2.49 μV, *SE* = 0.285) and out-of-the-zone periods (*M* = –2.42 μV, *SE* = 0.298) **(Figure S2B)**. The interaction between state and condition was also not significant, *F*(2,38) = 0.054, *p* = .948, η²p = .003, suggesting that selective attention is not modulated by the attentional states.

To further understand the neural mechanisms underlying lapse-prone states, we examined how inter-electrode correlations differed between correct and error trials as a function of attentional state. We computed inter-electrode correlations and contrasted correct vs. error trials separately for in-the-zone and out-of-the-zone periods. As shown in **Figure 3A and 3B**, the in-the-zone periods exhibited a widespread increase in inter-electrode correlations during correct responses compared to errors (FDR-corrected *p* < .05). In contrast, during out-of-the-zone periods, correlation differences between correct and error trials were markedly reduced, with fewer significant electrode pairs. These results suggest that periods of optimal attentional engagement (in-the-zone) are associated with enhanced large-scale synchrony of trial-evoked neural responses supporting successful performance, whereas such synchrony is diminished during lapse-prone states.

**Figure 3.**
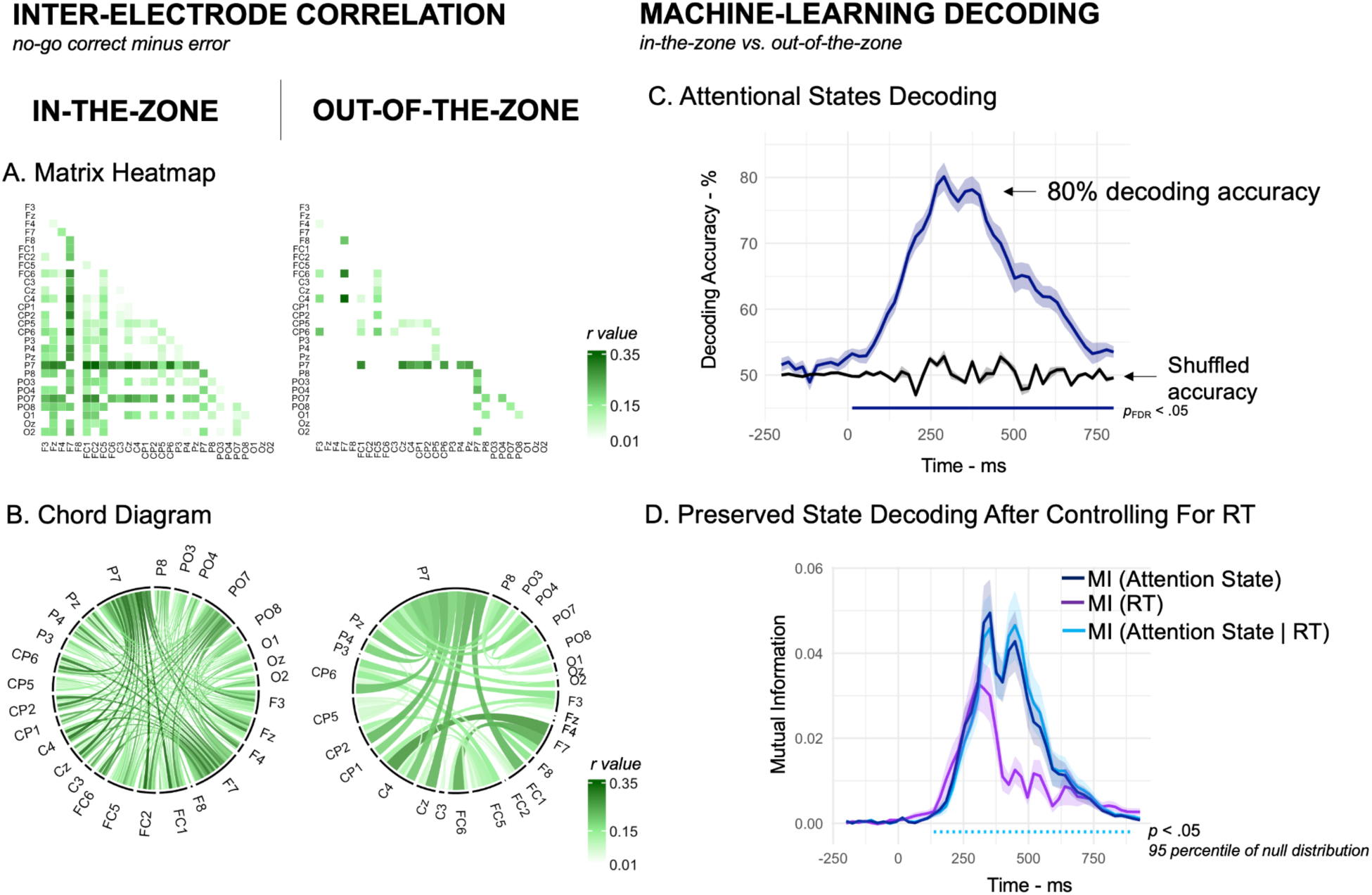
Neural signatures of lapse-prone states and their decoding with machine learning. The difference in inter-electrode correlation scores between correct and error no-go trials revealed increased correlation during in-the-zone periods, as shown in the correlation matrix **(A)** and the circular correlation diagram **(B)**. The classifier trained to detect attentional states from the EEG signal reaches 80% accuracy **(C)** and remains preserved after controlling for reaction time differences between the two states **(D)**. Shaded area represents the standard error of the group mean.

### Predicting Lapse-Prone States from EEG Signals

Now that the effect of poor sustained attention has been better characterized at the trial level and the state level, the second objective was to isolate unique brain signals predicting lapse-prone states. To this end, we trained a logistic regression-based machine learning classifier to decode attentional states from the EEG data. We focused our analyses on decoding sustained attention at the state-level—specifically, in-the-zone vs. out-of-the-zone periods—during correct go trials, which always involve a motor response. This approach allowed us to assess whether fluctuations in attentional state could be predicted independently of motor-related confounds. Our classifier was trained on 80% of the data and tested on the remaining 20%. This procedure was repeated 1,000 times, and performance metrics were averaged across iterations (*see SI Methods*).

Using this method, our classifier was able to decode participants’ sustained attention state with an accuracy reaching 80.1%. More precisely, a Wilcoxon signed-rank test on classification accuracy revealed a significantly higher decoding accuracy than the shuffled baseline starting at 13 ms and extending to 800 ms (all ps < .01, FDR-corrected). Peak accuracy reached 80.1% (SE = 2.15) at 289 ms, compared to a shuffled accuracy of 52.8% (SE = 0.96), with a Cohen’s *d* of 2.82 (Figure 3C).

Given the high decoding accuracy observed for distinguishing in-the-zone from out-of-the-zone states, we sought to ensure that this result was not merely driven by differences in reaction times or by systematic temporal patterns in the task sequence. To address this concern, we used a two-step analytical approach. First, we performed a partial mutual information analysis, estimating how much information EEG signals carried about attentional states, independently of RT. We computed the mutual information between the classifier’s trial-level predictions of attentional state, and the trial-level attention state labels defined by the VTC, while statistically controlling for RT, using a Gaussian copula estimation approach ^28^ (See SI Methods). This allowed us to isolate the unique contribution of brain signals to the prediction of attentional states, above and beyond behavioral variability. Second, to control for potential temporal confounds across nearby trials, we applied a circular shifting procedure: the labels for attentional states were randomly shifted across trials within each block, preserving the autocorrelation structure of the data while disrupting any fixed relationship between EEG and attentional states over time. The observed mutual information using the true labels was then compared to this null distribution. Together, these analyses confirmed that our classifier was genuinely decoding attentional state from EEG signals—not simply tracking RT fluctuations or trial timing—demonstrating that neural markers of lapse-prone states can be detected independently of behavioral or temporal cues **(Figure 3D).**

Once we identified a neural signal of attentional states, we sought to determine whether this signal is unique or whether it overlaps with selective attention or cognitive control. We therefore explored the extent to which the neural representation of attentional state differs from the attention allocated to task-relevant stimuli (left vs. right) and from the attention required to maintain the task set in mind (letter vs. number). To do so, we conducted Representational Similarity Analysis (RSA)—a multivariate method that examines how patterns of brain activity relate to cognitive variables of interest ^30,31^. Rather than simply decoding categories (i.e., in-the-zone vs. out-of-the-zone), RSA reveals how different task features shape the geometry of neural responses in representational space. To compute the neural dissimilarity matrix, we extracted the EEG voltage patterns across electrodes at each time point, treating the spatial configuration of brain activity as a multivariate feature vector. For each condition of interest, we computed the average topographical pattern and used a cross-validated Mahalanobis distance (crossnobis) to quantify dissimilarities between these patterns. This procedure was repeated across time to produce a time-resolved representational dissimilarity matrix (RDM) for each participant. We then compared a neural dissimilarity matrix to three theoretical dissimilarity matrices in which dissimilarity between the correct go trials is explained by attentional states (i.e., in-the-zone vs out-of-the-zone), selective attention (i.e., where the relevant stimulus appears—left or right), or cognitive control (i.e., which task the participant is performing—letter or number?) **(Figure 4A)** (*see SI Methods*). Averaged across time, all three predictors significantly explained variance in the representational structure: attentional states (mean semipartial *r* = 0.264, *p* < .001, VIF = 1.07), selective attention (*r* = 0.089, *p* = .007, VIF = 1.07), and cognitive control (*r* = 0.048, *p* = .022, VIF = 1.07) **(Figure 4A).** As all VIF values were close to 1, multicollinearity between regressors was minimal. An additional RSA included RTs as a nuisance regressor and showed that the state signal is also independent of RTs *(see SI Results),* supporting the results of the classifier. These findings show that neural representations were distinct between attentional states, selective attention and cognitive control, suggesting an independent EEG marker of lapse-prone periods.

**Figure 4.**
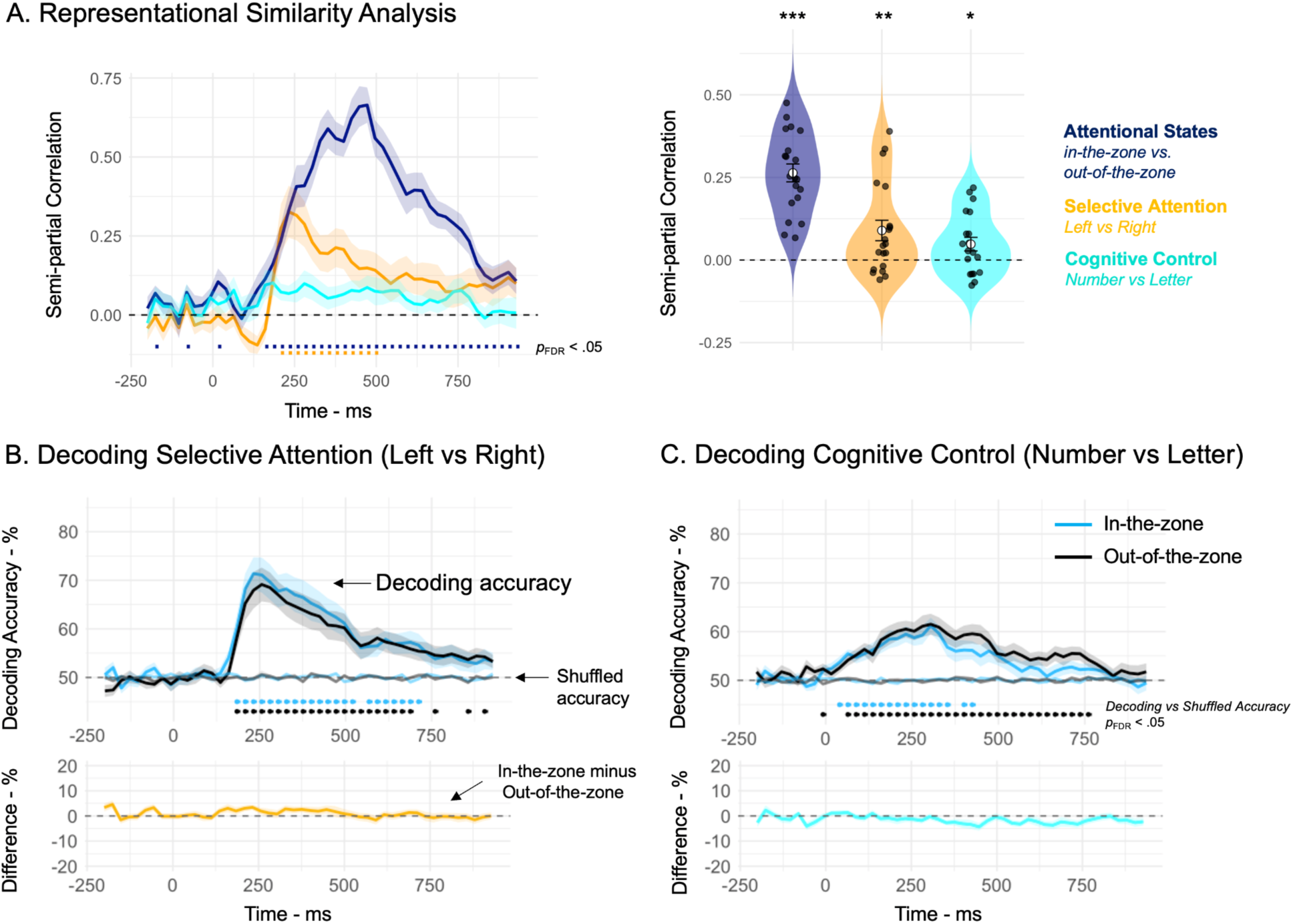
A Unique EEG Signal of Lapse-Prone States. Representational similarity analysis reveals that this EEG attentional state signal is distinct from both the selective attention and cognitive control **(A)**, suggesting a unique EEG signal of attentional states. Moreover, the classifier successfully decoded where participants directed their attention (left vs. right) **(B)** and which task they focused on (number vs. letter) **(C)**, performing significantly above shuffle accuracy. No differences in decoding accuracy were found between in-the-zone and out-of-the-zone periods. Shaded area represents the standard error of the group mean. \**p* < .05; \*\**p* < .01; \*\*\**p* < .001.

To take this a step further, we trained the classifier to decode selective attention during in-the-zone and out-of-the-zone periods. If attentional state impairs the ability to orient attention toward task-relevant stimuli (left vs. right), we would expect to observe a reduction in classification accuracy. A similar logic and analysis were applied to cognitive control (number vs. letter). The classifier successfully decoded selective attention, reaching 71% accuracy in-the-zone and 69% out-of-the-zone (**Figure 4B**), as well as cognitive control, achieving 61% accuracy for both in-the-zone and out-of-the-zone periods (**Figure 4C**). However, for both selective attention and cognitive control, the statistical analysis did not reveal any significant differences in decoding accuracy between in-the-zone and out-of-the-zone periods (*p_FDR_* > .05), suggesting that attentional states do not affect the ability to orient attention toward target stimuli or to maintain the task goal in mind.

## Discussion

The aim of this study was to better characterize the neural mechanisms modulated by periods of poor sustained attention and to test whether attentional states can be isolated from EEG signals beyond other attentional functions like selective attention or cognitive control. Our results showed that poor sustained attention at the trial level (also called lapses) and at the attentional state level (out-of-the-zone periods) was associated with reduced evoked brain responses, as revealed by ERP analyses and inter-electrode correlation measures. Furthermore, our machine learning analyses were able to decode a unique signal of attentional states, independently of behavioral variability and other attentional functions, with an accuracy reaching 80%.

Poor sustained attention at the trial level was studied through no-go errors, which are commonly referred to as attentional lapses in the literature. Our ERP analyses confirmed that these errors represent moments of lapse, as attention is withdrawn from the ongoing task, indicated by a reduction in the P300 component. This P300 reduction is similar to what has been observed during lapses in detection tasks ^14,27^. Additionally, and for the first time to our knowledge, our analyses revealed that attention lapses also impact the selection of task-relevant stimuli, as indicated by the reduced N2pc during lapses compared to correct no-go trials.

Poor sustained attention was also observed at the attentional state level, meaning not just on a single trial but over consecutive trials. To study this, we used the Esterman method to isolate “in-the-zone” periods (with low reaction time variability) and “out-of-the-zone” periods (with high variability) as markers of attentional states. In the bilateral CPT, we successfully isolated these periods, as shown by a higher percentage of no-go errors during out-of-the-zone periods compared to in-the-zone ^13,17,32,33^. ERP analyses also showed that attention was indeed withdrawn from the task during out-of-the-zone periods, as evidenced by a reduction in P300 amplitude compared to in-the-zone periods. Interestingly, unlike trial-level lapses, N2pc amplitude did not differ according to attentional state, suggesting that out-of-the-zone moments were not associated with impaired selection of task-relevant stimuli.

Beyond studying the neural mechanisms of poor sustained attention through specific electrodes, our inter-electrode correlation analysis aimed to better characterize the effect of sustained attention on global brain activity. We showed that poor sustained attention reduces the overall brain’s trial-evoked responses, both at the trial level and attentional state level. Specifically, inter-electrode correlations were stronger during correct no-go trials than during no-go error trials, indicating greater synchrony of trial-evoked responses during moments of sustained attention. Similarly, synchrony was higher during in-the-zone than out-of-the-zone periods, suggesting that lapse-prone states impact the coordination of the brain’s trial-evoked responses. This is consistent with previous studies showing that poor sustained attention reduces synchronization between brain regions ^34^ or between phases of neural oscillations ^13,15^.

After characterizing the neural mechanisms underlying sustained attention, we sought to test the feasibility of decoding lapse-prone states from EEG activity. We trained a classifier to detect in-the-zone vs. out-of-the-zone periods based on neural activity. The classifier achieved 80% accuracy. To ensure that the classifier was not merely decoding reaction time differences between attentional states, we conducted a mutual information analysis, which confirmed that the decoding reflected genuine attentional states beyond simple RT speed.

In an additional analysis, we showed that this attentional state signal is independent of other attentional functions, namely selective attention and cognitive control. Indeed, RSA analyses revealed that the state signal was distinct from EEG signals associated with attention allocated to the relevant stimulus side (left vs. right) or the relevant task-set (number vs. letter). Moreover, our classifier trained to decode selective attention and cognitive control showed no differences in accuracy between in-the-zone and out-of-the-zone periods, also suggesting that attentional state represents a distinct phenomenon that does not impair the efficiency of selective attention or cognitive control.

A follow-up question, then, is what underlies this unique EEG signal of lapse-prone states? One possible explanation proposed by Esterman and Rothlein (2019) is fluctuations in arousal level, although a recent study showed that participants do not report feeling sleepier during out-of-the-zone periods ^33^. An alternative explanation is opportunity cost. Out-of-the-zone state does not occur because one struggles to keep the goal in mind, but rather because other goals are more motivating than the current one. The unique EEG signal revealed by decoding could therefore be a measure of competition between goals, or of the effort required to pursue the current goal ^35^.

However, the decoding results are limited to VTC-based attentional states. Future studies could test whether selective attention and cognitive control are modulated when attentional states are measured using a subjective approach, such as mind-wandering reports, which capture a different form of attentional fluctuation ^32,33^. Similarly, decoding lapses at the trial level (i.e., errors) might be useful in different settings. As suggested by the observed reduction of N2pc amplitude during no-go errors, these brief lapses may represent a more extreme form of sustained attention failures where other attention functions may also be affected.

To conclude, by demonstrating the existence of a unique neural signal of sustained attentional states, this work carries both theoretical and practical implications. From a theoretical perspective, identifying an EEG marker of lapse would allow researchers to rethink the cognitive mechanisms underlying attentional fluctuations between attention states, during which cognitive control and selective attention appear to remain preserved. From a practical perspective, our results open the way to useful applications in everyday life. Real-time detection of poor sustained attention could be used to improve focus in schools, or to make driving safer by helping reduce the risk of road accidents.

## Materials and Methods

Materials and methods used in this study are detailed in SI Materials and Methods. Briefly, all participants provided written informed consent approved by the University of Chicago Institutional Review Board and were compensated for their participation. Participants completed a bilateral CPT (**Figure 1A**) lasting approximately 2 hours and consisting of 3,200 trials, allowing for a high-quality EEG signal-to-noise ratio. EEG data were continuously recorded during the task (250 Hz sampling rate, 0.01–80 Hz bandpass filter), and both electrophysiological and behavioral data were analyzed offline.

## Acknowledgments

We would like to warmly thank John Veillette for suggesting and discussing the mutual information analysis.

## Open practices

The data and codes are available on the Open Science Framework repository (https://osf.io/kw2fz/?view_only=5a26dd002b7a4523aaa5cf89c112f7ca)

## Author Contributions

Conceptualization: M.C., M.D.R., and E.K.V.; Data Curation: M.C.; Formal Analysis: M.C. and H.M.J.; Funding Acquisition: M.D.R. and E.K.V.; Investigation: M.C.; Methodology: M.C., H.M.J., M.D.R., and E.K.V.; Project Administration: M.C.; Resources: M.C., H.M.J., M.D.R., and E.K.V.; Software: M.C. and H.M.J.; Supervision: M.C., M.D.R., and E.K.V.; Visualization: M.C. and H.M.J.; Writing – Original Draft Preparation: M.C.; Writing – Review & Editing: M.C., H.M.J., M.D.R., and E.K.V.

## Supporting Information

### SI Methods

#### Participants

Twenty subjects participated in the study (mean age = 24.5, SD = 4.17, 10 females, 10 males). All participants reported normal or corrected-to-normal vision and normal color perception. Exclusion criteria were a history of neurological disorder. All participants gave their informed consent, and the protocol was approved by the local ethics committee. Subjects were paid for their participation.

#### Stimuli

Stimuli were presented on a gray background (90.0 cd/m²). Each trial began with the presentation of a red or blue square (3 × 3 cm) displayed at the center of the screen for 500 ms. This cue indicated whether participants should perform the letter task or the number task. Following the cue, a central black fixation point appeared for 300 ms. Then, two square images—one depicting a number and the other a letter (9 × 9 cm each)—were presented simultaneously for 600 ms, one to the left and one to the right of the fixation point. The image-side mapping of the two stimuli was randomly determined from trial to trial. The distance from the fixation point to the center of each image was 6 cm. After the stimulus offset, the black fixation point remained on the screen for 400 ms. It then turned green ([0 255 0]) for 600 ms, signaling to participants that they could blink if needed. Letters and numbers were generated using six different fonts, yielding 48 unique images for go trials and six different images for no-go trials. All images were randomly presented across trials.

#### Procedure

During the CPT, participants were instructed to categorize either letters or numbers in a go/no-go paradigm. If the cue color indicated the number task, participants were required to press a key for every number (go trials) except for the number 3, which served as the no-go stimulus. Conversely, if the cue color indicated the letter task, participants were instructed to press for every letter (go trials) except for the letter K (no-go trials). Cue colors were counterbalanced across participants. For each category, no-go trials occurred on approximately 15% of trials. The cue appeared every 20 trials, with the order of letter and number tasks randomized for each participant. Before beginning the main task, participants completed a short training session. The full task consisted of 3,200 trials and lasted approximately 2 hours. Participants were given three breaks during the task, each following the completion of 800 trials.

#### Behavioral and Statistical Analysis

The percentage of commission errors was defined as the number of errors on no-go trials divided by the total number of no-go trials. Mean reaction time (RT) was calculated based on correct responses on go trials. To assess attentional states, we computed intraindividual variability in RTs on correct go trials by calculating the Variance Time Course (VTC) for each participant. RTs were first normalized within subjects to standardize the VTC scale. The VTC was then derived for each trial as the absolute difference between that trial’s RT and the participant’s mean RT. For no-go error trials and trials without responses (e.g., omissions on go trials or correctly withheld responses on no-go trials), values were estimated via linear interpolation using RTs from adjacent trials. If a trial was preceded or followed by a cue or a break, we used the closest preceding or subsequent trial within the same block (i.e., trial *n–1* or *n+1*) to perform the interpolation. Performance was categorized into two attentional states based on the median VTC value across blocks: an in-the-zone state, characterized by reduced RT variability, and an out-of-the-zone state, characterized by increased RT variability. Paired *t*-tests were conducted to determine whether the percentage of no-go errors was significantly higher during out-of-the-zone periods compared to in-the-zone periods, with a two-tailed significance threshold of *p* < .05 .

#### EEG Recording

EEG activity was recorded using 30 active Ag/AgCl electrodes mounted in an elastic cap (actiCHamp, Brain Products, Munich, Germany). Electrodes were positioned according to the international 10–20 system at the following sites: Fp1, Fp2, F7, F3, Fz, F4, F8, FC5, FC1, FC2, FC6, C3, Cz, C4, CP5, CP1, CP2, CP6, P7, P3, Pz, P4, P8, PO7, PO3, PO4, PO8, O1, Oz, and O2. Two additional electrodes were placed on the left and right mastoids, and a ground electrode was positioned at Fpz. All electrodes were initially referenced online to the right mastoid and later re-referenced offline to the algebraic average of the left and right mastoids. Electrooculogram (EOG) activity was recorded using passive electrodes, with the ground placed on the left cheek. Horizontal EOG was measured using a bipolar configuration with electrodes placed approximately 1 cm from the outer canthus of each eye. Vertical EOG was recorded with a bipolar pair of electrodes placed above and below the right eye. Data were filtered online with a low cutoff of 0.01 Hz and a high cutoff of 80 Hz (12 dB/octave slope), and digitized at a sampling rate of 500 Hz using BrainVision Recorder (Brain Products, Munich, Germany) running on a PC. Electrode impedances were maintained below 7 kΩ during setup.

#### Eye-tracking

Gaze position was recorded using a desk-mounted infrared eye-tracking system (EyeLink 1000 Plus, SR Research, Ottawa, Ontario, Canada), with a sampling rate of 1,000 Hz. Participants maintained a stable head position throughout the task using a chin rest. The eye tracker was calibrated at the beginning of the experiment and recalibrated after each break.

#### Artifact Rejection

EEG data were bandpass filtered between 0.01 and 30 Hz using a second-order Butterworth filter. Artifact rejection was performed using an automated detection procedure, followed by manual visual inspection. Experimenters were blind to experimental conditions during artifact review. Trials containing artifacts were excluded from EEG analyses but retained for behavioral analyses. For saccades, trials were flagged as containing a saccade if the Euclidean distance between the mean gaze positions in the first and second halves of an 80-ms sliding window (advanced in 10-ms increments) exceeded 0.5° of visual angle. In the absence of usable eye-tracking data, horizontal EOG was used; trials were flagged if the mean voltage between halves of a 150-ms sliding window (advanced in 10-ms steps) differed by more than 20 μV. For blinks, trials were flagged as containing a blink if the eye tracker failed to detect the pupil at any time during the trial. When eye-tracking data were unavailable, vertical EOG was used; a blink was flagged when the mean voltage between the halves of a 150-ms sliding window exceeded 50 μV. For EEG artifacts, trials were flagged if the signal amplitude exceeded ±100 μV (absolute voltage threshold). Step-like changes were detected by identifying segments where the mean voltage differed by more than 60 μV between the first and second halves of a 250-ms sliding window, advanced in 20-ms steps. Linear drifts were flagged if the slope of a linear fit across the epoch exceeded 75 μV and the coefficient of determination (R²) was greater than 0.3. To identify high-frequency noise (e.g., muscle artifacts), trials were flagged when the peak-to-peak amplitude exceeded 75 μV within a 200-ms sliding window, advanced in 100-ms steps. Finally, trials were excluded if any channel flatlined for at least 200 ms within the rejection window. Known noisy electrodes, Fp1, and Fp2 were excluded from automated EEG artifact detection procedures to prevent them from biasing the rejection process and were also excluded from the analysis. On average, 24% of trials (SD = 15.1) were rejected from the analyses.

#### Event-related potential (ERP) analysis

EEG data were epoched from –200 ms to 950 ms relative to stimulus onset and baseline-corrected using the –200 to 0 ms pre-stimulus interval ^1^. The mean amplitude of each ERP component was computed within a ±50 ms window centered on the peak latency. The most negative peak for the N2pc was identified within the 200–400 ms post-stimulus window, and the most positive peak for the P3b was identified within the 300–700 ms post-stimulus window. The N2pc was calculated as the difference in mean activity between contralateral and ipsilateral electrodes relative to the location of task-relevant stimuli, focusing on electrodes P7 and P8, where the component was maximal. For the P3b, mean amplitude was extracted at electrode Pz, where the amplitude reached its maximum. Both peak amplitude and latency were extracted for each participant and condition.

#### ERPs statistical analysis

Statistical analyses were conducted at three distinct levels. First, to assess brain responses at the moment of a lapse, a three-factor repeated-measures ANOVA was performed on ERP data, comparing go correct, no-go correct, and no-go error trials. When significant effects were found, post-hoc *t*-tests were conducted, with Tukey’s correction applied for multiple comparisons. Second, to evaluate the effect of attentional state, a repeated-measures ANOVA trial type x states (in-the-zone vs. out-of-the-zone) was conducted on EEG amplitude.

#### Inter-electrode correlations

To investigate large-scale synchrony of trial-evoked neural responses during attention lapses, we conducted inter-electrode correlation analyses following the methodology of Hakim and colleagues ^2^. At the condition level, we computed for each participant the average EEG signal at each electrode separately for no-go correct and no-go error trials. We then calculated inter-electrode correlation matrices (Pearson’s *r*) from these condition-specific ERP time courses. This yielded one correlation matrix per condition and participant, quantifying the similarity of temporal activity profiles across all electrode pairs. Individual correlation matrices were then aggregated across participants. To statistically assess differences in inter-electrode correlations, we performed paired-sample *t*-tests at each electrode pair, comparing no-go correct and no-go error correlations across participants. This procedure yielded a full matrix of *t*-values and associated *p*-values. To control for multiple comparisons, we applied a false discovery rate (FDR) correction at *p* < .05 (Benjamini–Hochberg procedure), restricting statistical inferences to electrode pairs that survived this correction. The same procedure was applied to investigate interelectrode correlation during attentional states.

#### Multivariate Decoding Analysis

We implemented a logistic regression classification model using the same EEG segments employed for event-related potential analyses. To decode attentional states, we focused on correct go trials only. Due to their susceptibility to artifacts, electrodes Fp1 and Fp2 were excluded. To enhance the signal-to-noise ratio, trials were randomly grouped into sets of 20 without replacement within each attentional state, to improve classifier precision ^3^. Classification was conducted on averaged EEG activity using 48-ms time windows, progressing in 24-ms steps across the trial. At each time point, data were z-scored: training data were standardized, and testing data were normalized using the mean and standard deviation of the training set. Classifiers were trained on 80% of the data and tested on the remaining 20% using a stratified sampling approach to balance trials across the attentional states, along with the task-sets (letter vs number) and relevant target sides (left vs right). When the number of trial groups differed between conditions, the larger group was randomly down-sampled to match the size of the smaller one, thereby avoiding classification bias. This procedure was repeated 1,000 times, from the initial grouping through to testing, and performance metrics were averaged across iterations. Classification performance was evaluated using accuracy. To assess whether decoding performance exceeded chance level, we compared classification accuracy against an empirical baseline obtained through label shuffling. At each time point from stimulus onset, we performed a Wilcoxon signed-rank test across participants, comparing the observed accuracy to the shuffled accuracy. To control for multiple comparisons across time points, we applied a False Discovery Rate (FDR) correction using the Benjamini-Hochberg procedure with an alpha threshold of 0.05.

#### Mutual Information Analysis

Multivariate classification revealed strong decodability of attentional state. However, it is worth noting that participants were completing a go/no-go task involving frequent responses. These motor responses occurred within the trial epoch, producing response-related ERPs. In addition, attentional state was defined using the VTC, which is computed from reaction time. Thus, it could be that the decodability was driven by the classifier leveraging information about when exactly a motor response was going to occur on a given trial. To assess whether this is the case, we computed the partial mutual information (MI) between the EEG signal underlying classification and the state labels, while controlling for trial-level reaction times: MI(EEG; Attentional State | RT).

To isolate the EEG signal underlying classification, we first reran the classification analysis described above with the following changes. We ran a k-fold validation procedure with 10 folds, without any initial grouping or averaging of trials. After training on 90% of the data (with random down-sampling of the training set to balance the classification categories), each held-out trial’s distance from the classifier’s decision boundary was recorded. This was repeated with each fold serving as the test set once, so that each trial has a continuous classification value. The entire process was repeated 100 times with different initial folds, and each trial’s classification values were averaged across iterations.

We next compute the partial mutual information between the classification prediction values and the state labels defined by the VTC, while controlling for reaction times. To compute the mutual information, we applied the Gaussian copula estimation method ^4^ using the Frites package ^5^. This method can rapidly and accurately estimate mutual information between 2 variables, even when the relationship is non-linear.

The Frites package does not provide a direct method for computing the MI between one continuous variable and a discrete categorical variable while controlling for a different continuous variable, though it does have methods for computing the MI between 1) one continuous and one discrete variable, 2) two continuous variables, and 3) two continuous variables while controlling for a discrete variable. Thus, we computed the partial mutual information of interest via the following equation:

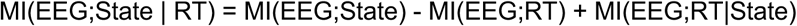

As MI is a positive value and cannot be compared against 0, we assessed its significance via permutation of the state labels. Because both state labels and reaction times may be autocorrelated across adjacent trials, we performed the permutation by circularly shifting the state labels by a random shift size for each block of each participant, then recomputing the MI of interest. Note that we performed the shifting before re-excluding trials marked with artifacts to preserve any autocorrelated structure across trials. To build up a null distribution, we repeated this permutation process 1000 times. To control for multiple comparisons across time, we took the largest MI value across time for each iteration, resulting in a null distribution of the 1000 largest MI values based on circularly shifted state labels. Timepoints were considered significant if they were greater than the 95th percentile of this distribution. These results were qualitatively identical if we performed the circular shifting within blocks, across blocks over the 800 trials within each run, or across the whole recording session of 3200 trials.

#### Representation Similarity Analysis

To investigate the structure of neural representations in our task, we conducted a representational similarity analysis (RSA) ^6,7^. RSA allows us to evaluate whether specific experimental factors are reflected in multivariate neural response patterns, under the assumption that cognitively similar conditions should elicit more similar neural activity. We focused on the same EEG segment used for ERP analysis and computed time-resolved representational dissimilarity matrices (RDMs) per subject using the cross-validated Mahalanobis distance (also known as crossnobis distance) ^8^. EEG data were first binned in sliding time windows of 48 ms with a 24 ms step, and RDMs were computed at each window across all channels (except Fp1 and Fp2). The RSA was performed on correct go trials, with the goal of assessing whether neural activity patterns reflect the attentional state (*in the zone* vs. *out of the zone*), task-set (letter vs. number) and stimulus side (left vs. right). Each condition was defined as a unique combination of these three factors. To estimate the crossnobis distance while avoiding overfitting, trials were stratified and randomly split into training and testing sets (50/50) across 1000 iterations. Within each fold, the number of trials was equalized across conditions to ensure balanced RDMs. Dissimilarities were computed between condition-specific means from train and test sets projected into a noise-normalized space using a Ledoit-Wolf regularized covariance matrix, estimated using the training set ^8^. The resulting subject-level RDMs were then averaged across permutations to obtain a single time-resolved RDM per subject. For each RSA, we defined theoretical RDMs corresponding to the expected dissimilarity between condition pairs based on the factorial structure of the task. Theoretical RDMs were computed by taking the absolute difference between condition values: for categorical factors, the dissimilarity was 0 when the conditions matched and 1 otherwise. For each subject and time point, empirical RDMs were regressed against the theoretical RDMs using rank-based multiple linear regression ^9^. This yielded semipartial correlations for each regressor, reflecting its unique contribution to the neural similarity structure. Collinearity between regressors was assessed using the Variance Inflation Factor (VIF), and semipartial correlations were tested at each time point using Wilcoxon signed-rank tests. False Discovery Rate (FDR) correction (Benjamini-Hochberg) was applied to correct for multiple comparisons across time points.

### SI Results

#### Performance in the letter *versus*. the number task

To verify that the two tasks are similar in difficulty, a paired-samples t-test comparing mean reaction times between the number (M = 386.3 ms, SD = 41.2) and letter tasks (M = 386.1 ms, SD = 40.0) showed no significant difference, *t*(19) = 0.08, *p* = .937, Cohen’s *d* = 0.02. A second paired-samples t-test on no-go errors between the number and letter tasks revealed no significant difference between the number task (M = 25.5, SD = 14.8) compared to the letter task (M = 28.0, SD = 15.2), *t*(19) = –1.92, *p* = .070, Cohen’s *d* = –0.43).

#### N2pc

To explore whether the amplitude difference between correct and error no-go trials is due to a modulation in the processing of either the relevant or the irrelevant side, an ANOVA was conducted on contralateral and ipsilateral amplitudes. On the contralateral amplitude, the ANOVA revealed a significant main effect of trial type, *F*(2,38) = 23.0, *p* < .001, η²p = .547. Post-hoc t-tests showed significantly greater N2 amplitude for no-go correct compared to go trials, *t*(19) = 5.60, *p* < .001, p_tukey_ < .001; and greater for no-go correct compared to no-go error trials, *t*(19) = –5.13, *p* < .001, p_tukey_ < .001. The difference between go and no-go error trials was not significant, *t*(19) = 1.70, *p* = .106, p_tukey_ = .232. The ANOVA performed on ipsilateral N2 amplitude revealed no significant main effect of trial type, *F*(2,38) = 0.02, *p* = .981, η²p = .001 (**Figure S1**).

**Figure S1.**
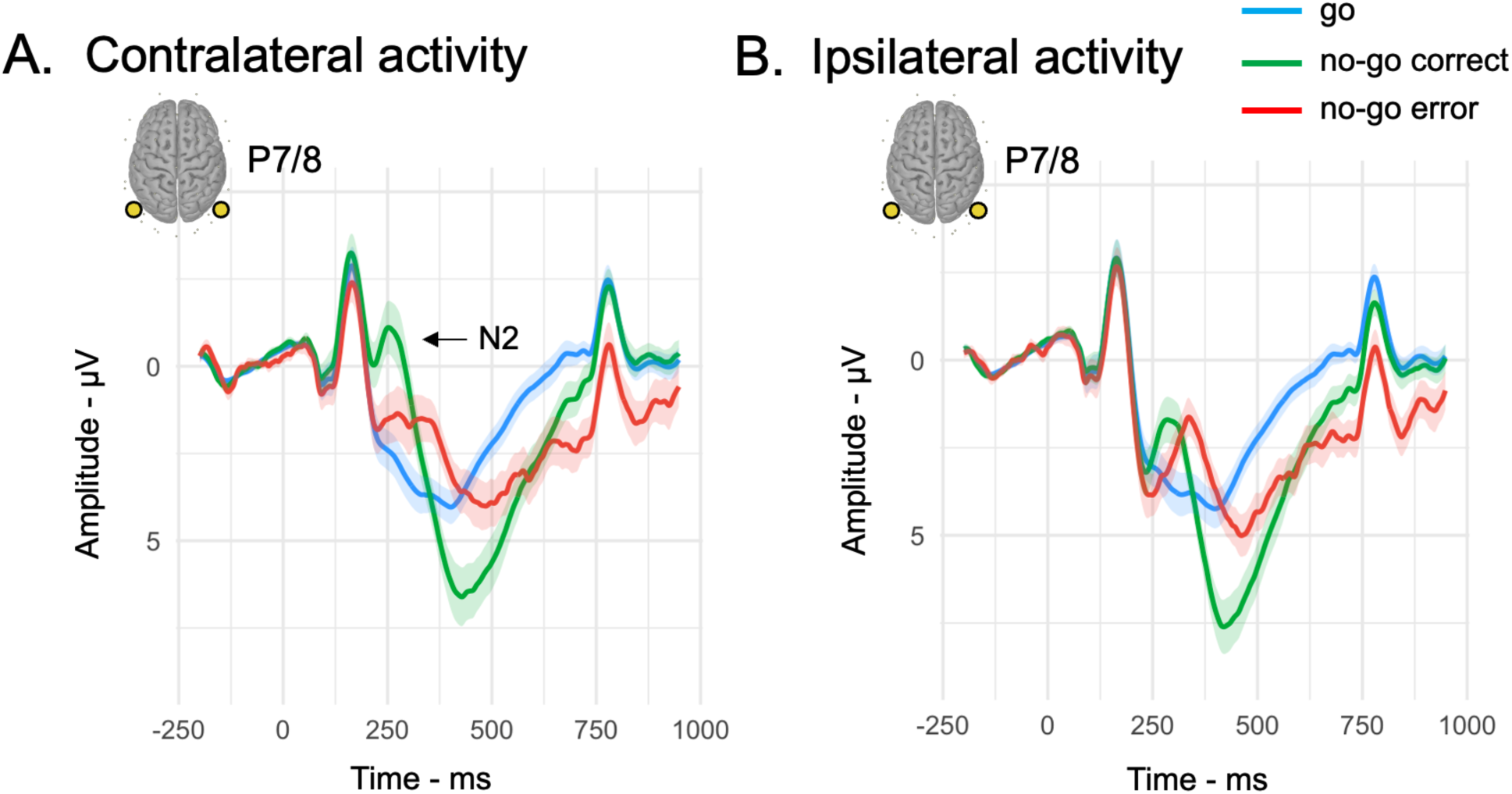
Lateralized N2 during attentional lapses. Reduced N2 amplitude on contralateral electrodes during sustained attention lapses (A). N2 amplitude on ipsilateral electrodes does not differ between trial types (B). Shaded area represents the standard error of the group mean.

#### RSA

By adding the RT factor, attentional state (mean semipartial *r* = 0.257, *p* < .001, VIF = 1.10) and relevant side (*r* = 0.089, *p* = .006, VIF = 1.07) explained significant variance in the representational structure. In contrast, neither reaction time (*r* = 0.037, *p* = .131, VIF = 1.04) nor task type (*r* = 0.042, *p* = .057, VIF = 1.09) reached statistical significance. These findings suggest that neural similarity patterns were primarily driven by fluctuations in attentional state and spatial features, rather than reaction speed or goal-relevant task parameters. All variance inflation factors (VIFs) were close to 1, indicating negligible collinearity among regressors.

#### Decoding selective attention and cognitive control during attentional states

We trained the classifier to decode selective attention (left vs. right) during in-the-zone and out-of-the-zone periods, as well as cognitive control (number vs. letter). The parameters and statistical tests used were the same as those previously applied to decode attentional states. The classifier successfully decoded selective attention, reaching 71% accuracy in-the-zone and 69% out-of-the-zone (**Figure 4B**), as well as cognitive control, achieving 61% accuracy for both in-the-zone and out-of-the-zone periods (**Figure 4C**). Although the periods of successful classification were similar for selective attention between in-the-zone and out-of-the-zone, the decoding of the task-set was sustained for a longer duration out-of-the-zone (up to ∼760 ms after stimulus onset) compared to in-the-zone (up to ∼425 ms after stimulus onset). This suggests that maintaining the task-set requires more sustained neural engagement out-of-the-zone compared to in-the-zone.

**Figure S2.**
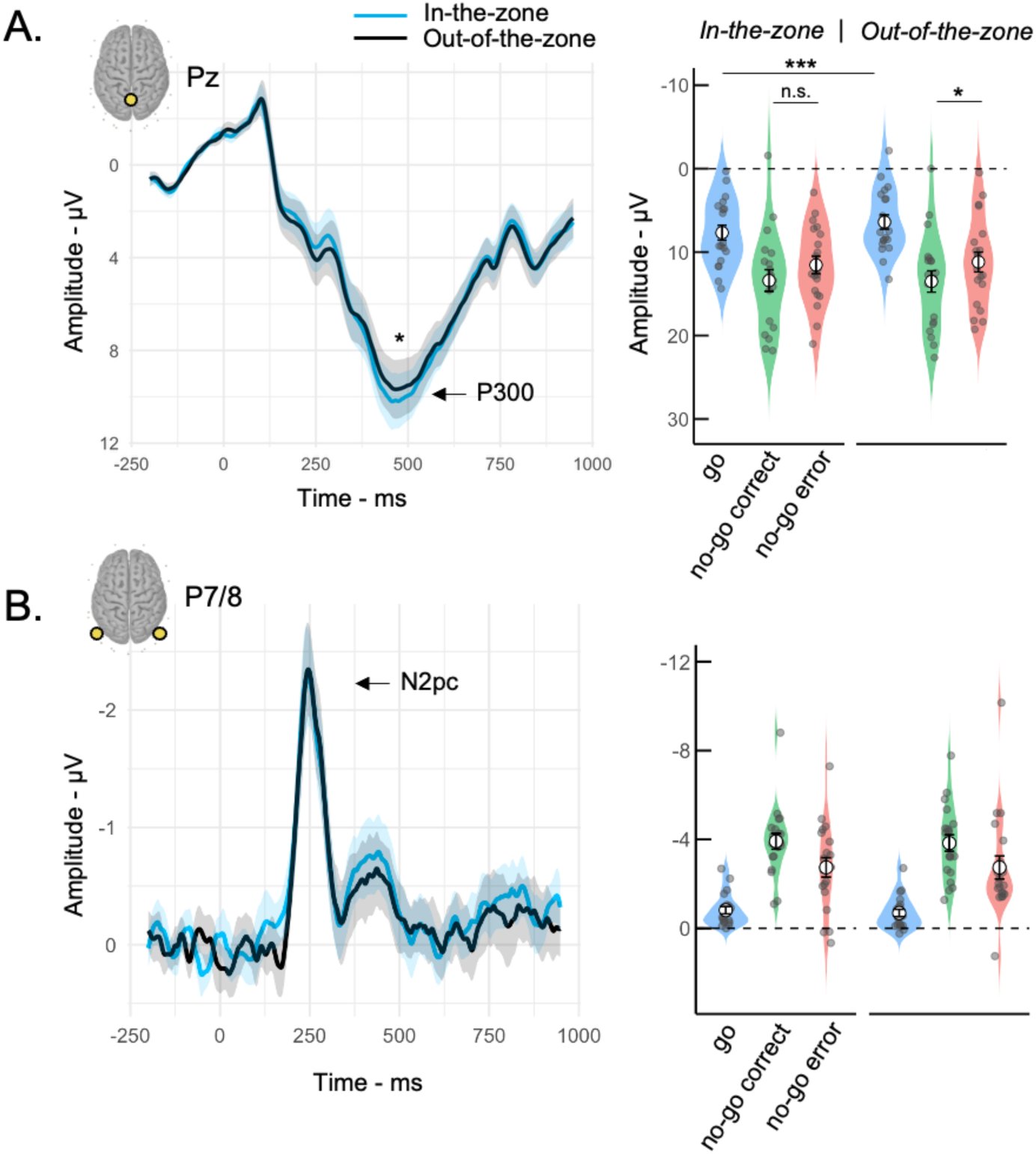
Neural mechanisms underlying attentional states. Lapse-prone state was assessed through the out-of-the-zone periods or periods of high RT variability. The amplitude of the P300 was reduced out-of-the-zone compared to in-the-zone states (A), but the N2pc amplitude was not modulated by attentional states (B). Shaded area represents the standard error of the group mean.

